# Dynamic role of GlyT1 as glycine sink or source: pharmacological implications for the gain control of NMDA receptors

**DOI:** 10.1101/2024.03.17.585409

**Authors:** Stéphane Supplisson

**Affiliations:** Institut de Biologie de l’ENS (IBENS), Ecole normale supérieure, Université PSL, CNRS, INSERM, Paris, F-75005, France

**Keywords:** GlyT1, NMDAR, Glycine transporter, sarcosine, heteroexchange, coagonist

## Abstract

Glycine transporter 1 (GlyT1) mediates termination of inhibitory glycinergic receptors signaling in the spinal cord and brainstem, and is also diffusely present in the forebrain. Here, it regulates the ambient glycine concentration influencing the ‘glycine’-site occupancy of *N* -methyl-d-aspartate (NMDARs). GlyT1 is a reversible transporter with a substantial, but not excessive, sodium-motive force for uphill transport. This study examines its potential role as a glycine source, either by reversed-uptake or by heteroexchange. I explored how glycine accumulation triggers its release, facilitating the activation of NMDARs by glutamate applied alone. Indeed, glutamate evokes no current in “naive” oocytes coexpressing GluN1/GluN2A and GlyT1, a previously characterized cellular model, but now using GlyT1 as the only potential source of coagonist for NMDAR activation. After glycine uptake, however, glutamate evokes large currents, blocked by ALX-5407 and potentiated by sarcosine, a specific inhibitor and substrate of GlyT1, respectively. These results suggest higher occupancy of the co-agonist site when GlyT1 functions as a glycine source either by reversed-uptake or by heteroexchange. A difference between these two glycine-release mechanisms occurs at hyperpolarized potentials, which induce an apparent voltage-dependent block of NMDAR currents, whereas heteroexchange preserves NMDAR activation at these potentials. Together, these results confirm GlyT1-mediated efflux as a positive regulator of NMDAR co-agonist site occupancy, and demonstrate sarcosine heteroexchange effectiveness in enhancing coagonist site occupancy. Depending on its actual mode of transport, GlyT1-inhibitors and sarcosine may have distinct effects on ambient glycine and NMDAR facilitation, and be a source of variation in reversing NMDAR hypofunction in schizophrenia.

**Graphical Abstract:** 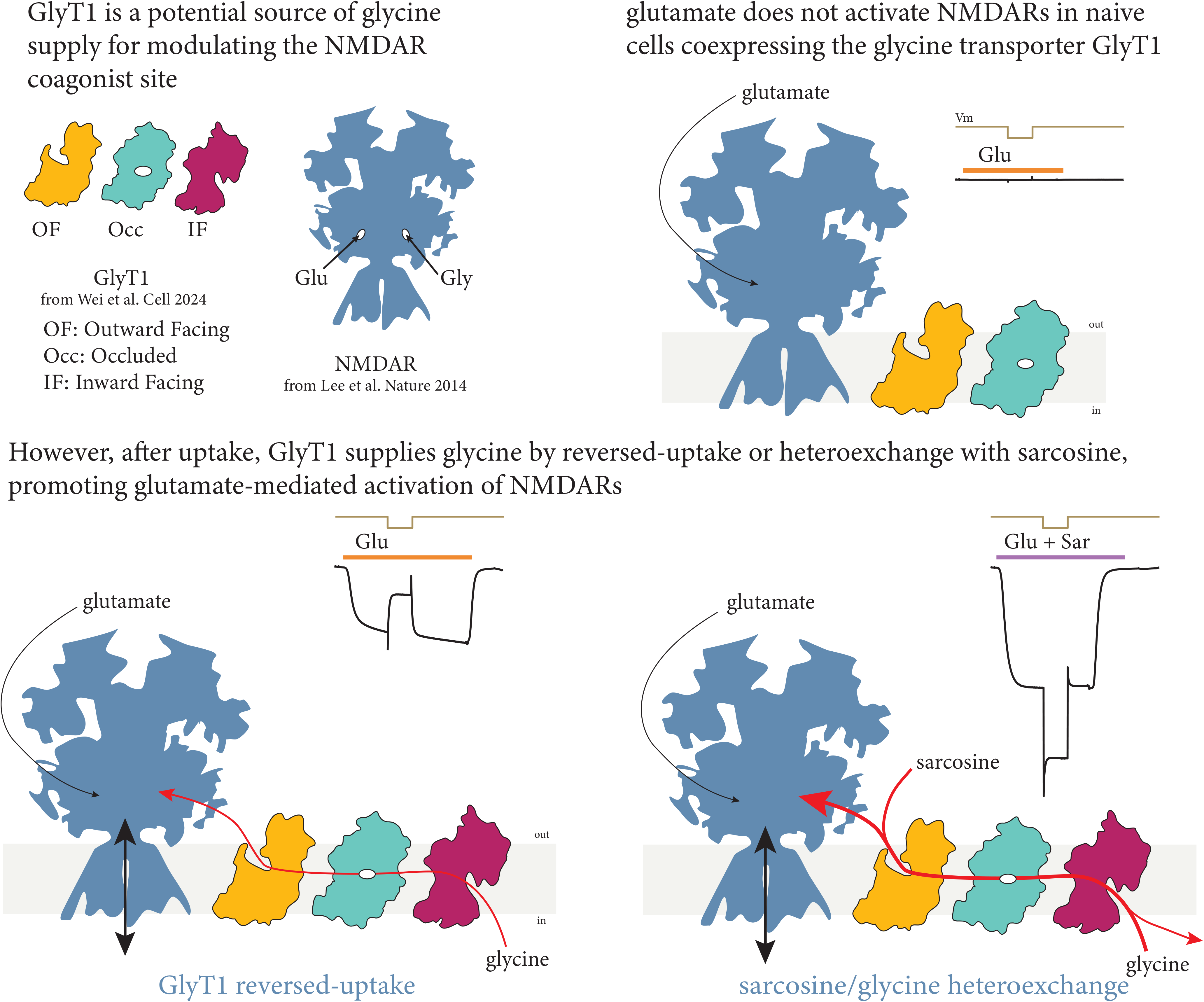

## 1. INTRODUCTION

Glycine acts as a high-affinity co-agonist for NMDA receptors at excitatory synapses (Johnson and Ascher, 1987). Its binding to the GluN1 subunit enables their activation by glutamate in heterologous expression system (Kleckner and Dingledine (1988), see recent reviews in Hansen et al. (2021); Mony and Paoletti (2023)). This seminal discovery by Johnson and Ascher revealed that glycine is spontaneously released by primary cultures of cortical and diencephalic neurons (Johnson and Ascher, 1987), suggesting that a glycine transporter, likely the glial isoform GlyT1, operates in reversed uptake mode in the absence of glycine (Attwell et al., 1993; Shibasaki et al., 2017).

To explore how GlyT1 switches from glycine sink to glycine source function and modulates co-agonist site occupancy, I recorded currents in GluN1/GluN2A- and GlyT1-coexpressing oocytes, a simple diffusion-limited cellular model described previously (Supplisson and Bergman, 1997; Supplisson and Roux, 2002; Guellec et al., 2022). Heteromeric assembly of GluN1/GluN2A subunits forms ‘classical’ (i.e. glutamate- and glycine-gated) NMDARs with the lowest sensitivity for glycine (Paoletti et al., 2013), requiring much larger glycine supply for activation than high-affinity GluN1/GluN2B receptors (Guellec et al., 2022).

### GlyT1 uptake desaturates NMDAR co-agonist site

GlyT1 belongs to a small cluster of four amino-acid transporters in the solute carrier 6 (SLC6A9) family (Bröer and Gether, 2012). Its structure has been solved in three conformations (outward-facing, occluded and inward-facing) with typical LeuT-fold of SLC6 transporters (Yamashita et al., 2005; Kristensen et al., 2011; Shahsavar et al., 2021; Wei et al., 2024).

In agrement with a 2:1:1 Na^+^:Cl^−^:glycine stoichiometry, glycine uptake is electrogenic, carrying 1 elementary charge per glycine; therefore the time integral of the transport current is directly proportional to glycine accumulation (Roux and Supplisson, 2000).

In the CNS, GlyT1 is primarily found in fine astrocytic processes and subset of glutamatergic terminals (Zafra et al., 1995; Cubelos et al., 2005), and plays a crucial role in maintaining ambient glycine levels below saturation for NMDARs (Smith et al., 1992; Attwell and Bouvier, 1992; Supplisson and Bergman, 1997; Bergeron et al., 1998; Supplisson and Roux, 2002; Zafra et al., 2017; Piniella and Zafra, 2023). Its main function, however, is to recapture glycine released at inhibitory synapses in the spinal cord, brainstem, cerebellum and retina (Zafra et al., 1995; Gomeza et al., 2003; Eulenburg et al., 2018; Eulenburg and Hülsmann, 2022). As iGlyRs have much lower sensitivity to glycine compared to NMDARs (Legendre, 2001; Paoletti et al., 2013), GlyT1 termination of inhibitory signaling allows crosstalk with excitatory synapses, and NMDAR facilitation by spillover of glycine released at these sites has been demonstrated in the dorsal horn of the spinal cord (Ahmadi et al., 2003).

High glycine concentrations, exceeding 100 µM in bath application, are necessary to overcome GlyT1 uptake and potentiate NMDARs in brainstem slices (Berger et al., 1998). However, experiments using glycine-puffs through pressure ejection or light-uncaging of CNI-glycine uncovered the activation of GluN1/GluN3A receptors, which form atypical excitatory glycine-gated receptors (eGlyR) in the medial habenula (Otsu et al., 2019) and amygdala (Bossi et al., 2022, 2023). Moreover, evidence that d-serine can substitute for glycine in the forebrain, further suggests that GlyT1 reduces below saturation the ambiant glycine sensed by NMDARs (Papouin et al., 2012; Bail et al., 2015; Ferreira et al., 2017).

GlyT1 is a drug target for schizophrenia based on the hypothesis linking NMDAR hypofunction to this psychiatric disorder (Javitt, 2012; Umbricht et al., 2014; Uno and Coyle, 2019). Different classes of compounds have been developed that inhibit selectively GlyT1 (Atkinson et al., 2001; Aubrey and Vandenberg, 2001; Harvey and Yee, 2013; Hofmann et al., 2016; Al-Khrasani et al., 2019; Rosenbrock et al., 2023), and elevate ambient glycine in hippocampal brain slices and *in vivo*, enhancing NMDARs and causing a small but tonic inhibition by activating iGlyRs (Bergeron et al., 1998; Chen et al., 2003; Zhang et al., 2008; Sipilä et al., 2014). In contrast, competitive inhibition of glycine uptake by sarcosine (N-methylglycine, a GlyT1-specific substrate (Supplisson and Bergman, 1997; Vandenberg et al., 2007; Guellec et al., 2022)) could in addition enhance glycine efflux by heteroexchange as described below (Herdon et al., 2001; Hanuska et al., 2016).

### GlyT1 reversed uptake and heteroexchange promote glycine efflux

Despite its high sodium-motive force favoring glycine uptake (Attwell et al. (1993), see appendix in Guellec et al. (2022)), GlyT1 must reverse in the absence of external glycine, since there is no influx to balance efflux. However, there is almost 200-fold difference between external (23 µM) and internal (4.3 mM) glycine EC_50_, estimated in GlyT1-expressing oocytes and CHO cells, respectively (Roux and Supplisson, 2000; Aubrey et al., 2005). Consequently, basal glycine efflux is negligible in naive GlyT1-expressing oocytes, which contain low, sub-millimolar glycine concentrations (Taylor and Smith, 1987; Meier et al., 2002); in agreement, there was no facilitation of GluN1/GluN2B receptors in naive oocytes expressing GlyT1 (Guellec et al., 2022). Studies using tracers, as well as electrophysiological recordings, have shown evidence for trans-stimulation of glycine efflux by homoexchange or heteroexchange with sarcosine (Zafra and Gimenez, 1986; Zafra and Giménez, 1988; Huang et al., 2004; Aubrey et al., 2005; Guellec et al., 2022).

### Diffusion-limited unstirred layer in Xenopus oocyte

*Xenopus* oocytes are large cells, with millimeter diameter, mechanically protected by a fibrous vitelline membrane (Wolf et al., 1976). Their plasma membrane forms numerous invagination and microvilli that expand their surface by ∼7-fold, creating an unstirred layer limiting the exchange time even in high-flow chambers (Barry and Diamond, 1984; Costa et al., 1994; Supplisson and Bergman, 1997). Reduction of passive diffusion in this extracellular unconvected fluid compartement (Occhipinti et al., 2014), is an advantage for studying how transporters deplete or accumulate their substrate in a small, not well stirred, juxtamembrane environment. Ultimately, creating a stopped-flow condition that limits diffusion of the bath solution to the membrane enhances the juxta membrane depletion of the substrate resulting from its uptake (Supplisson and Bergman, 1997; Zuo and Fang, 2005; Sun et al., 2014; Sheipouri et al., 2020).

## 2. EXPERIMENTAL PROCEDURES

### Coexpression of NMDAR GluN1-GluN2A and GlyT1 in Xenopus oocytes

Capped mRNAs were synthesized using the Ambion mMessage mMachine T7 Transcription Kit (Thermo Fisher Scientific) from cDNAs encoding rat GlyT1b and NMDAR subunits GluN1 and GluN2A. These cDNAs were subcloned in a modified pRC/CMV vector, as previously described in Supplisson and Bergman (1997). Oocytes were harvested by gentle shaking of saclike ovary isolated from female *Xenopus lævis* (TEFOR Paris-Saclay, CNRS/INRAE, Université Paris-Saclay) in Ca^2+^-free OR-2 solution (in mM: 85 NaCl, 1 MgCl_2_, 5 Hepes, pH 7.6 with KOH) containing 5 mg ml^−1^ collagenase (type A, ROCHE). Defolliculated oocytes were injected with 50 ng mRNAs of mixed GlyT1, GluN1, and GluN2A mRNAs using a nanoliter injector (World Precision Instruments). Post-injection, oocytes were maintained at 19 °C in individual wells filled with 1 ml of Barth’s solution (in mM: 88 NaCl, 1 KCl, 0.41 CaCl_2_, 0.82 MgSO_4_, 2.5 NaHCO_3_, 0.33 CaNO_3_, 5 Hepes, pH 7.4 (with NaOH)) containing 50 µg*/*ml gentamycin and 50 µM d-APV.

### Electrophysiology

Three to eight days post-injection, two-electrode voltage-clamp (TECV) recordings were performed using an OC-725D amplifier (Warner Instruments) with oocytes coexpressing GlyT1 and NMDAR. Before experiments, oocytes were preincubated in 30 µM BAPTA-AM for 20 to 30 min to help buffer intracellular Ca^2+^ during long recordings. Glass electrodes filled with 3 M KCl solution had a typical resistance of 1 MΩ. Oocytes were held at −40 mV, and perfused continuously with a solution containing (in mM) 100 NaCl, 1.8 CaCl_2_, 1 MgCl_2_, 5 Hepes, pH 7.2 (with KOH) at 25 °C. Glutamate was applied in Ca^2+^- and Mg^2+^-free Ringer solution (replaced by 0.3 mM BaCl_2_). As low divalent increases the leakage current, Ba^2+^ Ringer was applied only 10 s before and maintained 15 s after each glutamate application. Glycine uptake was performed in normal CaCl_2_ and MgCl_2_ Ringer containing 10 µM d-APV. Currents were typically filtered at 100 Hz and acquired at 1 kHz using a Digidata 1440A and the pCLAMP 10 software suite (Molecular Devices).

Weighted time constants 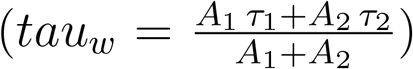 were calculated from the amplitude and time constant of the exponential decay time course during washout.

Glycine accumulation was derived from the time-integral of the uptake current as: 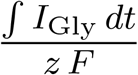, with *z* represents GlyT1 charge coupling (1.01 e/gly, Roux and Supplisson (2000); Vandenberg et al. (2007)), and *F* is the Faraday’s constant (96 485 C mol^−1^).

### Chemicals

Salt, buffer and amino acids were obtained from Merck| Sigma-Aldrich. Chemicals and drugs were obtained from TOCRIS (d-APV, ALX-5407) and Merck| Sigma-Aldrich (ALX-5407, BAPTA-AM)

## 3. RESULTS

This study examines how GlyT1-mediated glycine efflux facilitates NMDAR activity using two-electrode voltage clamp recordings on *Xenopus* oocytes expressing only GluN1/GluN2A (GlyT1^−^) or co-expressing these with GlyT1 (GlyT1^+^).

### GlyT1 functions as an ambient glycine sink controlling NMDAR activation

In GlyT1^−^ oocytes, coapplication of glutamate and glycine evoked NMDAR currents of amplitude comparable to or slightly greater than those recorded with d-serine as a co-agonist (2 µM each, Figure 1A). In contrast, GlyT1^+^-oocytes showed a notable reduction in ambient glycine levels, as the current amplitude decreased by 54 % compared to the response in d-serine (from 112 *±* 9 % (n = 7) to 51.6 *±* 6.3 % (n = 9), Figures 1B, 1C). This effect is even larger when the membrane is stepped at a more negative potentials, which increase the driving force for both, NMDAR currents and GlyT1 uptake. Increased uptake results in further glycine depletion, leading to an immediate increase in current amplitude that is followed by relaxation to a steady-state, but surprisingly with little or no change in amplitude (Figure 1B). One minute application of ALX-5407 (5 µM) inhibited GlyT1 and restored the glycine current full amplitude (120 *±* 12 %, n = 4), higher that induced by d-serine (Figures 1B,1C). Notably, GlyT1 appears to accelerate the rate of glycine clearance during its washout, as indicated by a reduced weighted decay time constant (tau_*w*_). In GlyT1^+^ oocytes, the mean tau_*w*_ was 487 *±* 58 ms (n = 8), which is 44 % lower than the tau_*w*_ calculated for oocytes lacking transporter binding sites for buffering and capturing glycine (1105 *±* 111 ms, n = 22, P = 6.42 *×* 10^−5^ Wilcoxon rank-sum test; this second group encompasses tau_*w*_ from GlyT1^−^-oocytes, application of d-serine, and inhibition by ALX-5407, as their means were not statistically different, P = 0.78).

**Figure 1:**
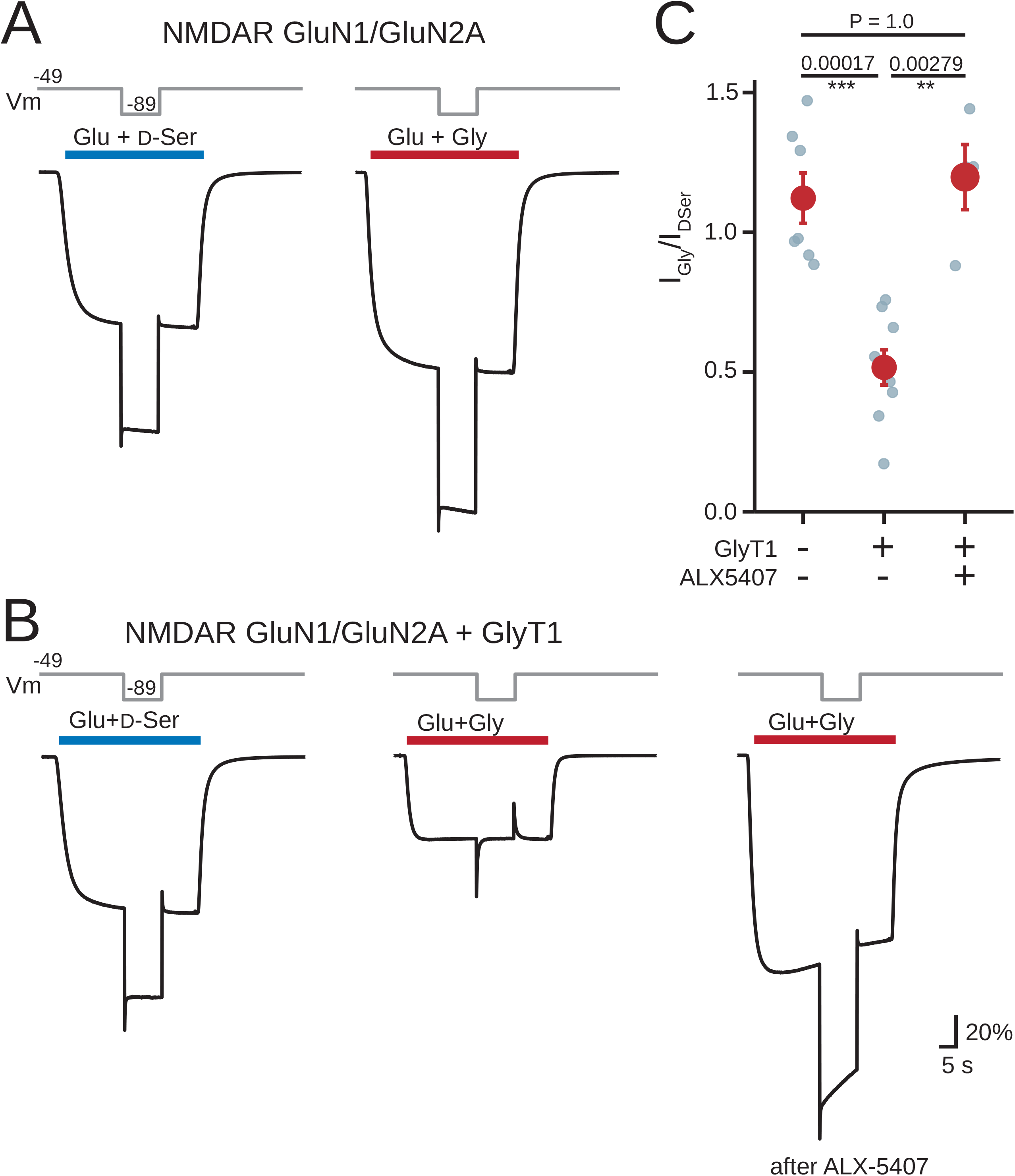
GlyT1 decreases ambient glycine sensed by the NMDAR co-agonist site in oocytes coexpressing GluN1/GluN2A subunits. **(A)** In control (GlyT1^−^) oocytes expressing GluN1/GluN2A, currents evoked by glutamate (Glu) in combination with d-serine (d-Ser) or glycine (Gly) exhibits similar amplitudes, which increase upon a negative voltage step. All co-agonists were applied at 2 µM. **(B)** In GlyT1^+^ oocytes, glycine uptake reduces NMDAR activation compared to d-serine. Moreover, a negative voltage step does not increase the current amplitude at steady-state. Inhibition of GlyT1 after one minute application of 5 µM ALX-5407 restored normal glycine response. **(C)** Summary of GluN1/GluN2A currents evoked by 2 µM glutamate + glycine in GlyT1^−^, and GlyT1^+^ oocytes in control and after one minute incubation with 5 µM ALX-5407. Current amplitudes were normalized to the absolute amplitude of the current evoked with 2 µM d-serine. Error bars indicate SEM, with n the number of oocytes; two-side unpaired Wilcoxon rank-sum test: ***P < 0.001, **P < 0.01, *P < 0.05.

### GlyT1 functions as a glycine source facilitating NMDAR activation

#### Glycine supply by GlyT1-mediated reversed-uptake

Preliminary stopped-flow experiments were conducted to provide initial evidence of NMDAR activation facilitated by GlyT1-reversal. In GlyT1^+^ oocytes exhibiting low transporter expression, the stopped-flow condition accentuates the reduction in [Gly]_*jm*_ thereby decreasing co-agonist site occupancy and NMDAR current, whereas interrupted flow has no effect when d-serine served as co-agonist (Figure 2A).

**Figure 2:**
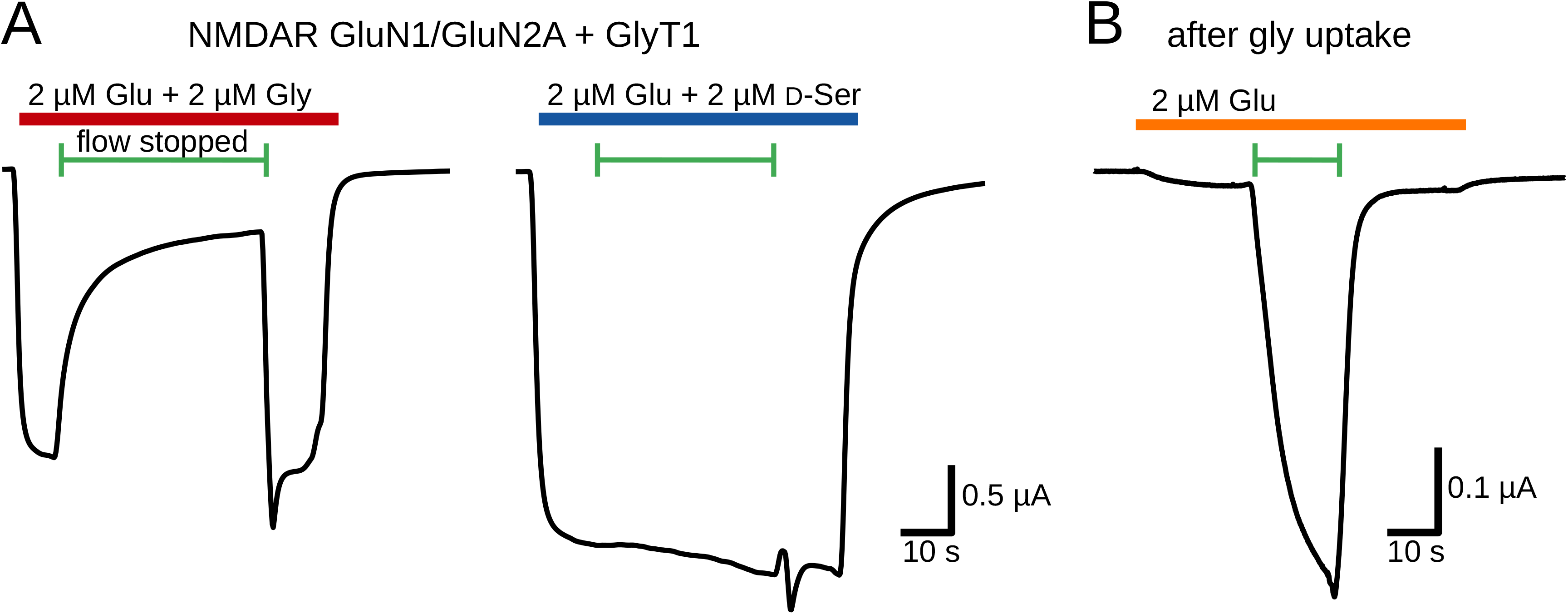
Stopped-flow condition enhances glycine depletion or extracellular accumulation induced by GlyT1. **(A)** Interrupting the flow of a solution containing 2 µM glutamate and glycine enhances the reduction in glycine concentration sensed by the co-agonist site in oocytes coexpressing GluN1/GluN2A and GlyT1 (left), whereas it has no significant effect when d-serine is used as co-agonist (right). **(B)** After 3 min uptake with 200 µM glycine (in normal Mg^2+^, Ca^2+^ ringer and 10 µM d-APV), application of 2 µM glutamate alone evokes a small NMDAR current suggesting some glycine supply by GlyT1 reversed-uptake. Arrest of the flow triggered a massive (x 28-fold) increase in NMDAR current, suggesting extracellular accumulation of ambient glycine under stopped-flow condition.

Figure 2B presents a typical recording involving a low expressing GlyT1^+^ oocyte after glycine uptake followed by an extended period of washout (*I*_*Gly*_ =−33 nA for 200 µM glycine in normal Mg^2+^- and Ca^2+^-Ringer with 10 µM d-APV; the time-integral of the glycine-evoked current was 13.9 µC for 3 min application). Application of glutamate alone evoked a modest NMDAR current (−17.7 nA), but its amplitude increases 28-fold (to −496 nA) under stopped-flow condition, suggesting that a small glycine efflux by GlyT1 reversed-uptake generated a larger accumulation of ambient glycine in the stopped-flow condition, enhancing co-agonist site occupancy (Figure 2B).

These glycine accumulations were often unstable and susceptible to perfusion artifacts in the open chamber used in this study. Therefore efflux experiments were performed in continuous flow using GlyT1^+^ oocytes with high transporter expression (Figure 3). For each oocyte tested, applications of glutamate (2 µM), alone or with sarcosine (50, 100, 200 µM) were repeated for three distinct and successive conditions defined as: 1) **naive**, as initially, oocytes have low intracellular stock of glycine since they are maintained for several days in individual wells containing 1 ml of Barth solution supplemented with gentamycin and d-APV; 2) **after glycine uptake** (applying 200 µM glycine for 3 to 15 minutes); 3) **after ALX-5407** (one minute application of 5 µM ALX-5407 induces an “irreversible” inhibition of GlyT1) Current amplitudes were normalized to the response evoked by glutamate and d-serine in naive oocytes (2 µM each).

**Figure 3:**
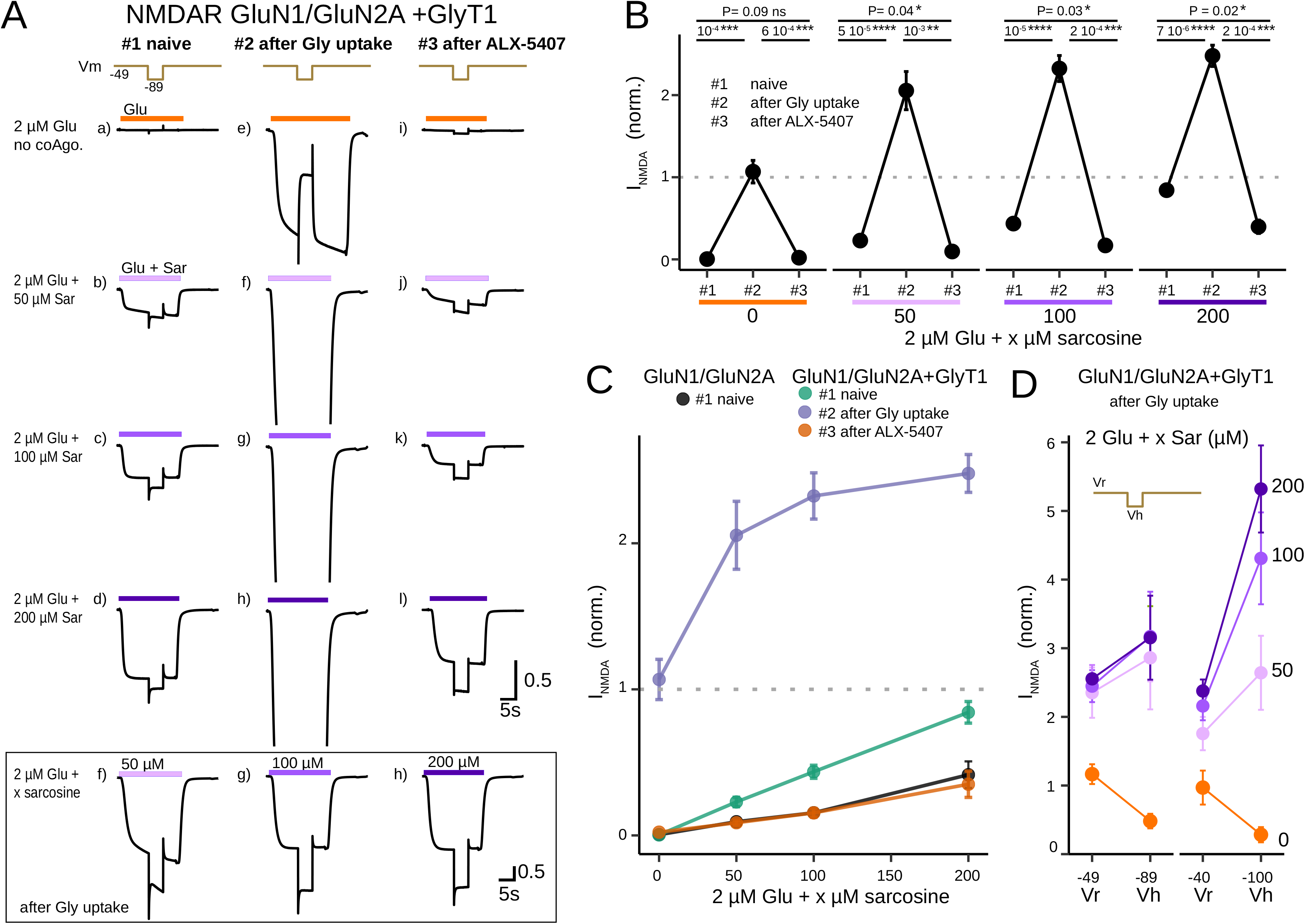
The supply of glycine by GlyT1, via reversed-uptake or heteroexchange with sarcosine, facilitates glutamate-gated activation of NMDARs. GlyT1^+^-oocytes coexpressing GluN1/GluN2A were selected for high transporter expression, with uptake current amplitude larger than −50 nA for 200 µM glycine at −40 mV. **(A)** Array of 15 current traces [a-l] recorded with the same oocyte for applications of 2 µM glutamate alone [a,e,i] or with sarcosine (50 µM [b,f,j], 100 µM[c,g,k] or 200 µM [d,h,l]). Each application was repeated for 3 distinct conditions which are organized in columns for: #1) naive oocytes (left column, [a-d]); #2) after glycine uptake (middle column, [e-h]; time integral of the uptake current: 86.5 µC); #3) after 5 µM ALX-5407 (right column, [i-l]; one minute incubation). The bottom panel shows the unclipped-traces for sarcosine after glycine uptake [traces f-h]. **(B)** Summary of the current amplitude change for the three conditions (naive, after Gly uptake, after ALX-5407) for each application (no coagonist, 50 µM Sar, 100 µM Sar, 200 µM Sar). Paired t-test, n = 5 for each application. **(C)** Sarcosine dose-response for application of 2 µM Glu as function of the sarcosine concentration for the three conditions (naive (green), after uptake (purple), after ALX-5407 (red)) for GlyT1^+^ oocyte. The black curve is the same protocol for naive GlyT1^−^ oocytes. Currents were normalized by the amplitude of the response with 2 µM d-Ser as co-agonist, as shown with a dotted line. **(D)** Opposite voltage dependency of NMDA current for hyperpolarized potential for glycine supply by reversed uptake (x=0, orange) or heteroexchange with sarcosine (50, 100 and 200 µM, purple). Two different voltage steps were applied (−40 mV and −60 mV) from different holding potentials, as indicated in the axis legend.

In naive GlyT1^+^ oocytes, glutamate alone failed to evoke a current, confirming that GlyT1 is not a glycine source *per se* under basal condition (Figures 3Aa, 3B, 3C). However, after uptake, GlyT1 reversal provides a continuous glycine supply that modulates the co-agonist site occupancy, thus enabling glutamate-applied-alone signaling, with current amplitude comparable to those evoked with 2 µM d-serine (Figures 3Ae,3B). Larger glycine accumulation were achieved in high-expressing oocytes, as shown by a mean time-integral of the uptake current of 80.4 *±* 5.7 µC (n = 7), corresponding to 833 *±* 58 pmol of glycine (mean uptake current: −186 *±* 39 nA (n = 7, from −322 to −79 nA at −40 mV); glycine application was 200 µM for 8.1 *±* 1.5 min at an average Vm of −80 mV).

A hyperpolarizing voltage step during glutamate application increased the driving force for NMDAR current while reducing glycine efflux, resulting in a higher initial current that relaxed to a lower steady-state amplitude (Figures 3Ae, 3B, and 3D), demonstrating GlyT1 indirect control of NMDAR activation. Inhibiting GlyT1 with ALX-5407 (5 µM) blocked the only source of extracellular glycine and terminated the facilitation of NMDAR activation by glutamate alone (Figures 3Ai, 3B). These findings collectively demonstrate that glycine efflux, by GlyT1 reversed-uptake in glycine-loaded oocytes, facilitates glutamate-gating of low-affinity GluN1/GluN2A receptors effectively (Figures 3Ae, 3B, and 3C).

#### Glycine supply by sarcosine heteroexchange facilitates NMDAR activation

Sarcosine is a weak NMDAR co-agonist (McBain et al. (1989), but see Zhang et al. (2009)). In GlyT1^−^ oocytes, coapplication of 2 µM glutamate and 200 µM sarcosine induced small currents, with amplitude 41.5 *±* 9.1 % (n = 7) of the response with 2 µM d-serine as a co-agonist. In naive GlyT1^+^ oocytes, however, the glutamate + sarcosine current represents 84.2 *±* 7.4 % (n = 8, P=0.0033, Welch Two Sample t-test) of the glutamate + d-serine response. This potentiation suggests that sarcosine may already facilitate glycine supply by heteroexchange, whereas glycine efflux was not detected by reversed-uptake in this condition (see above). It should be noted, however, that part of the current induced by 200 µM sarcosine is due to its electrogenic uptake as a GlyT1 substrate, although the precise contribution of this transport current was not quantified. Indeed, after glycine uptake, potentiation of the NMDAR current by sarcosine far exceeds this initial difference, indicating that GlyT1-mediated heteroexchange greatly increases glycine efflux and NMDAR co-agonist site occupancy compared to reversed-uptake condition. All traces in Figure 3A are plotted with the same scale, resulting in the clipping of the three sarcosine traces after uptake. These three traces are presented below with a different current scale for clarity.

The potentiation of NMDAR current by 50 µM sarcosine (896 %, from 0.229 *±* 0.032 (n = 9) to 2.05 *±* 0.23 (n = 8), P=8.924 *×* 10^−5^, Welch Two Sample t-test) appears to saturate at higher concentrations (Figure 3C), reflecting either saturation of glycine efflux and/or high occupancy of the co-agonist site. As expected, GlyT1 inhibition by ALX-5407 reduces the glutamate-gated current to levels comparable to those recorded in GlyT1^−^ oocytes (Figure 3C). Additionally, membrane hyperpolarization with a voltage step does not affect heteroexchange as the glutamate current amplitude increases, unlike the reduction observed with reversed uptake, a contrast highlighted in the summary Figure 3D.

#### GlyT1-block of NMDAR-facilitation by reversed uptake at negative potentials

To examine these differences in the voltage dependence of glycine supply by GlyT1 reversed-uptake or heteroexchange, current-voltage (I-V) relationships were constructed for different conditions as shown in Figure 4A. The I-V curves, generated in response to 2 µM glutamate either applied alone (glycine is provided by reversed uptake, left panel) or coapplied with 50 µM sarcosine (glycine is provided by heteroexchange, right panel), were constructed using slow voltage ramps under the three conditions previously defined for GlyT1^+^ oocyte (naive, post-glycine uptake, and after ALX-5407 inhibition). Notably, the I-V curve for glycine supply via reversed uptake (shown as blue curve in the left panel in Figure 4A) exhibits a pronounced voltage-dependent block with a negative slope, in apparent similarity with the Mg^2+^-block of NMDAR (Nowak et al., 1984). However, these recordings were conducted in a 0.3 mM Ba^2+^ Ringer solution, without Mg^2+^ and Ca^2+^, suggesting that this voltage-dependent inhibition must be indirect, probably caused by a decrease in co-agonistsite occupancy as glycine relase reduces at negative potentials. Indeed, sodium-ions are transported against their electrochemical gradients during a complete reversed-uptake cycle, as illustrated in the schema of Figure 4B (left panel), thus limiting the rate of glycine efflux at negative voltages. In contrast, the I-V curve for sarcosine heteroexchange after glycine uptake (blue curve in the right panel of Figure 4A) appears almost linear, displaying only minor deviation at very negative potentials as NMDAR becomes sensitive to traces of Mg^2+^ (Kuner and Schoepfer, 1996; Retchless et al., 2012). This linear I-V supports that sarcosine/glycine heteroexchange is electroneutral, as it involves no net transfer of the ions cosubstrates (Figure 4B). These I-Vs are consistent with the observed difference in current amplitude during hyperpolarized voltage steps, as described in Figure 3D. In naive oocytes, the flat I-Vs (orange curves) confirm negligible glycine efflux via reversed uptake, except for Vm*>*+20 mV. Furthermore, the presence of an NMDAR outward current in sarcosine supports that heteroexchange with glycine occurs even with low intracellular glycine levels in naive oocytes. Finally, the absence of glutamate-evoked current after incubation with 5 µM ALX-5407 confirms the lack of glycine release by endogenous amino acid transporters of *Xenopus* oocytes.

**Figure 4:**
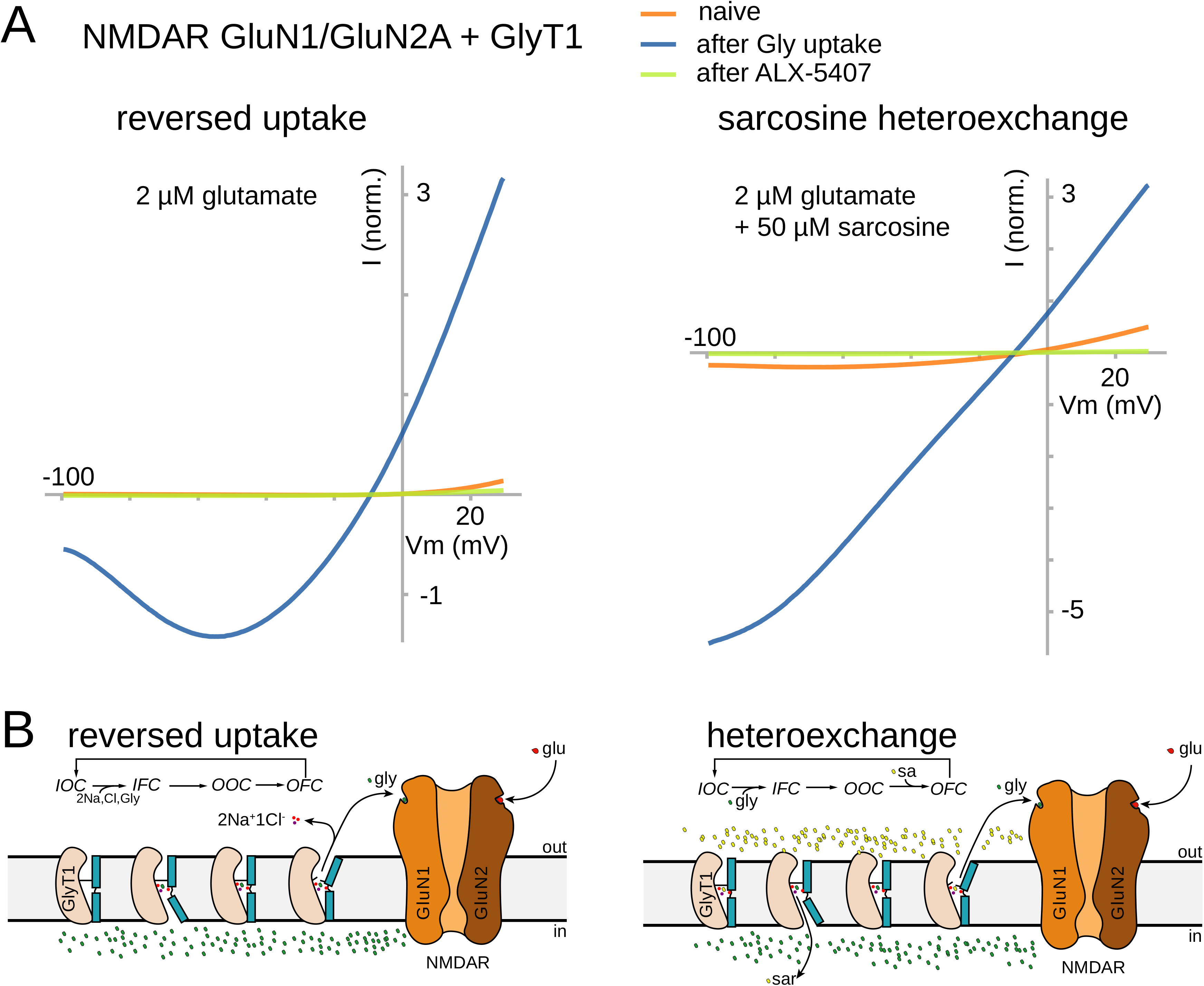
Current-voltage (I-V) relationships of glutamate-evoked NMDAR currents in GlyT1^+^ oocyte depending on the mode of glycine supply. **(A)** Representative I-V curves were obtained from the same GlyT1^+^ oocyte under three different conditions: naive, post-glycine uptake, and after 5 µM ALX-5407. Currents were recorded by applying 10-second voltage-ramps from −100 to +30 mV before and during application of 2 µM glutamate either alone (left panel) or with 50 µM sarcosine (right panel). Net evoked current were obtained by subtraction. As [Gly]_*jm*_ change with voltage-dependent reversed-uptake by GlyT1, the use of a slow voltage-ramp protocol (13 µV ms^−1^) allows to approach dynamic equilibrium for [Gly]_*jm*_ and NMDAR activation at each potential. Currents were normalized to the response elicited by 2 µM d-serine as co-agonist. **(B)** Schemetic diagrams illustrate the GlyT1 transport cycle for glycine efflux via reversed-uptake (left) and sarcosine heteroexchange (right), based on the structural dynamics of SLC6 transporters, which support an alternative access mechanism of transport. GlyT1 is represented in four conformations using the Rocking model (Drew and Boudker, 2015; Fan et al., 2021) as template: Outward Facing (OFC), Inward Facing (IFC), Outward Occluded (OOC) and Inward Occluded (IOC). Each step is reversible, but not mentioned in the schema for simplicity.

#### The decay time-course of NMDARs facilitation by GlyT1 reversed-uptake depends on their sensitivity to glycine

After several minutes of uptake and glycine subsequent washout, a steady outward current is recorded in GlyT1^+^ oocytes, indicating an electrogenic glycine efflux by GlyT1-mediated reversed-uptake (see Figure 4B). The average outward current (+17.4 *±* 2.3 pA, n = 7) represents 9.4 % of the uptake current recorded at near saturating glycine concentration (200 µM, −186 *±* 39 pA, n = 7). Consequently, glycine accumulations produced by uptake are transient overshoots that decrease over time as glycine is released.

To explore the decay time course of NMDAR facilitation, glutamate-evoked currents were recorded at different time points before and after glycine uptake in GlyT1^+^ oocyte (Figure 5A). While facilitation of NMDAR activation greatly amplifies the signal generated by glycine efflux, it also distorts the decay kinetics due to the narrow range of sensitivity to glycine. A decay time t_1*/*2_ of 9.8 min was measured in the experiment shown in Figure 5A, indicating that tonic glycine supply allows glutamate-gating of NMDAR for a long period post-uptake. This slow decay timecourse is even more pronounced with GlyT1^+^ oocytes coexpressing high affinity GluN1/GluN2B receptors, as previously shown (Figure 5B is adapted from supplementary Figure 16 in Guellec et al. (2022)), Remarkably, the amplitude of the glutamate-evoked current remained at 6.5 µA, i.e. 80% of the initial value, one hour after glycine washout.

**Figure 5:**
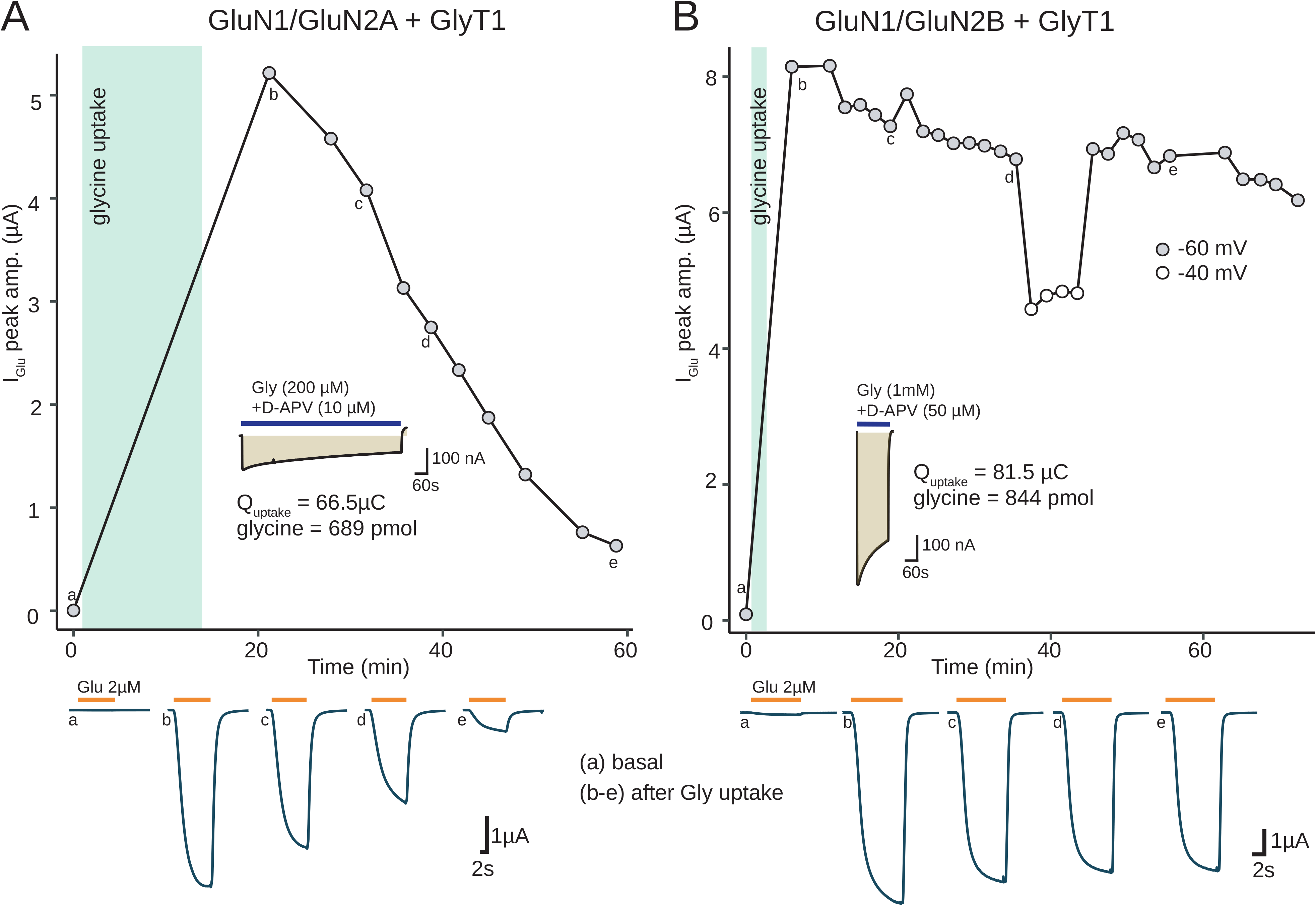
Post-uptake, GlyT1 maintains glycine supply and high occupancy of the NMDAR co-agonist site; the decay time of potentiation depends on GluN2 subunit sensitivity to glycine. **(A-B)** Peak amplitude of the current evoked by 2 µM glutamate before (a) and after (c-e) glycine uptake in GlyT1^+^ oocytes coexpressing either (A) GluN1/GluN2A or (B) GluN1/GluN2B. The current traces below correspond to different time points before and after glycine uptake, as indicated by a letter. The GluN1/GluN2B panel is adapted from the supplementary Figure 16 in Guellec et al. (2022). The uptake currents are displayed with the same current and time scales in both panels, so surface integrals are comparable.

## 4. DISCUSSION

### GlyT1 reversed-uptake as a glycine source for NMDAR co-agonist site

This study demonstrates, in a coexpression model, that GlyT1 can function as a steady source of glycine supply to modulate NMDAR co-agonist site occupancy.

It is posited that GlyT1 can deplete or accumulate glycine within a small juxtamembrane compartment ([Gly]_*jm*_), which is sensed by the NMDAR co-agonist site (Supplisson and Bergman, 1997). Under constant perfusion, [Gly]_*jm*_ dynamics depend on two fluxes: the GlyT1-mediated net flux across the membrane (*J*_Gly_) and passive diffusion (*J*_dif_) though an unstirred layer between the bath and the juxtamembrane compartment.

*J*_Gly_ is directly proportional to the transport current (*I*_Gly_):

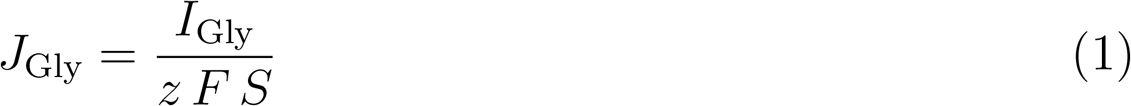

where *z* and *F* have already been defined and *S* represents the spheric surface of the ooocyte (0.038 cm^2^ for a diameter of 1.1 mm).

According to Fick’s first law of diffusion, the glycine flux (*J*_dif_), between the bath compartment ([Gly]_*b*_) and the juxtamembrane compartment is given by :

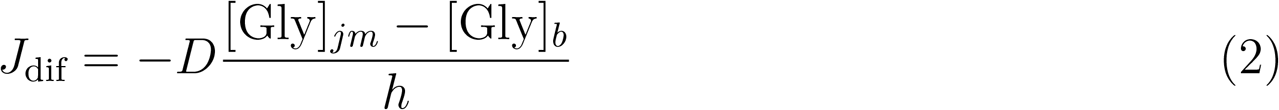

where *D* is the effective coefficient of diffusion for glycine (5*×*10^−6^ cm^2^*/*s), and *h* is the thickness of the unstirred layer. When oocytes are perfused in a glycine-free solution ([Gly]_*b*_ = 0), the diffusional flux simplifies to:

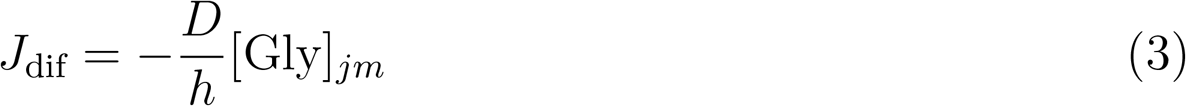

A steady-state condition is reached when *J*_dif_ = −*J*_Gly_, for which [Gly]_*jm*_ detected by the NMDAR co-agonist site corresponds to:

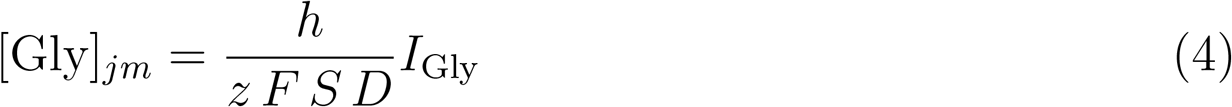

This equation predicts stationnary [Gly]_*jm*_ between 1 and 1.9 µM for an average outward current of *I*_Gly_ = +17 nA, assuming a range of 11 to 20 µm for *h* (Costa et al., 1994; Supplisson and Bergman, 1997).

These values confirm GlyT1’s capability to modulate ambient glycine concentrations within a range suitable for GluN1/GluN2A-receptors (Figure 3), comparable to the response evoked by 2 µM d-serine (or glycine) as co-agonist.

### Efficacy of sarcosine in facilitating glycine release by heteroexchange

Previous reports have shown that sarcosine stimulates glycine heteroexchange in GlyT1-expressing cells (Herdon et al., 2001; Huang et al., 2004; Hanuska et al., 2016). This trans-stimulation of glycine efflux, also observed for glycine itself (Zafra and Giménez, 1988; Guellec et al., 2022), produces a futile cycle that reduced GlyT1 uptake current as glycine accumulates in the cell (Roux and Supplisson, 2000; Aubrey et al., 2005).

Although it may seem counterintuitive that GlyT1, a tightly coupled cotransporter, would exhibit this futile cycle of exchange process, where influx of one substrate facilitates efflux of another, evidence under voltage-clamp conditions confirms this trans-stimulation. Specifically, external glycine triggers an inward current in GlyT1-expressing oocytes — indicating a net glycine influx— while simultaneously increasing the unidirectional efflux of ^14^C-glycine, a trans-stimulation not observed in GlyT2-expressing oocytes (Guellec et al., 2022).

In the absence of external glycine, the ion-motive force provided to GlyT1 by the coupling of 2 Na^+^ and 1 Cl^−^ remains intact, directed inwards. However, reversed-uptake is a complete backward-transport-cycle requiring the dissociation of all three ions from the outward-facing conformation and their re-association in the inward-facing conformation (Figure 4B). Both steps are unfavorable at the holding potential of −40 mV given the inwardly-directed electrochemical gradients for Na^+^. Consequently, more hyperpolarized potentials that reduce glycine efflux exponentially (Roux and Supplisson, 2000), cause indirectly a voltage-block of NMDAR current due to reduced glycine occupancy of the co-agonist site.

In contrast, sarcosine driven heteroexchange is electroneutral as it bypass the requirement to form an empty transporter in the Inward Occluded conformation (Figure 4B). Therefore the heteroexchange transport cycle is not influenced by voltage and benefits from two favorable substrate gradients: inwardly-directed for sarcosine and outwardly-directed for glycine.

Since the unidirectional glycine efflux triggers by sarcosine heteroexchange is not proportional to the transport current, a kinetic model of the GlyT1 transport cycle is needed to estimate glycine release by this mechanism, although, it may be difficult to infer transporter kinetics from an indirect readout of glycine efflux as detected here with NMDAR activation, due to the narrow sensitivity range of the coagonist site.

Sequential and random order of binding models have been proposed for GlyT1 alternative access cycle, but exchange remains largely unexplored (Aubrey et al., 2005; Erdem et al., 2019). Findings from these models differ as they predict either a higher rate of reverse translocation (Aubrey et al., 2005), or alternatively, high allosteric cooperativity between cosubstratres sites, which is specific of GlyT1, that facilitates the reverse mode and glycine release in a symmetrical model (Erdem et al., 2019).

In contrast, amphetamines heteroexchange of dopamine or serotonine have been studied in detail for the monoamines transporters DAT and SERT, with different mechanisms, models, and transporter structures (Jones et al., 1999; Schicker et al., 2012; Wang et al., 2015; Sitte and Freissmuth, 2015; Hasenhuetl et al., 2018; Navratna and Gouaux, 2019). Fast reverse mode and exchange has also been characterized for excitatory amino acid transporters (EAATs) — albeit coupled to a symport of 3 Na^+^ and 1 H^+^, and antiport of 1 K^+^, and with a different transport mechanism than GlyT1 —, and structures of several intermediate states of EAAT symport/antiport cycle are now documented (Kavanaugh et al., 1996; Zhang et al., 2007; Canul‐Tec et al., 2022; Qiu and Boudker, 2023).

### Post-uptake, glycine accumulation triggers a steady overflow and NMDAR facilitation

To switch on glutamate signaling in GlyT1^+^ oocytes, glycine must first accumulate significantly by uptake. This intracellular build-up leads to a tonic overflow of glycine by GlyT1 reversed-uptake after external clearance, increasing coagonist site occupancy and facilitating glutamate-gated activtation of NMDAR. The mean time-integral of the uptake current (80 µC) predicts glycine accumulation of 833 pmol, a value that exceeds by 9 to 14 times the amount of glycine measured in freshly dissociated oocytes (ranging from 59 to 92 pmol (Taylor and Smith, 1987; Meier et al., 2002)). Naive oocytes, which have been deprived of external amino acids for days, may have even lower basal glycine levels. Assuming an oocyte (stage 5-6) water volume of 0.4 ml (Taylor and Smith, 1987), the estimated cytosolic glycine concentration would rise from 147 to 230 µM for naive oocytes to ∼ 2.1 to 2.3 mM after uptake. These ranges of concentration are in good agreement with the EC_50_ (4.3 mM) estimated for GlyT1 outward current in mammalian CHO cells (Aubrey et al., 2005), suggesting an efficient but non-saturating glycine supply after uptake. In contrast, negligible glycine efflux is expected in naive (untreated) oocytes, in agreement with the absence of response to glutamate. Interestingly, the mean glycine concentration after uptake, corresponds to the ∼ 2 mM estimate for the concentration of glycine in astrocytes proposed by Attwell et al. (1993), based on Berger et al. (1977).

### Sarcosine as a glycine releasing agent

One motivation of this study was to clarify the mode of action of sarcosine, often described as a GlyT1 inhibitor or glycine reuptake inhibitor, and used as a tool to effectively increase ambient glycine levels, like in the amygdala (Bossi et al., 2022), whereas NFPS, a “true” GlyT1 inhibitor, produced a slow accumulation in the same brain structure (Li et al., 2013). However, the results obtained with this coexpression model demonstrate that in the absence of extracellular glycine, thus ruling out inhibition of uptake, sarcosine is indeed a potent glycine-releasing agent by GlyT1-mediated heteroexchange when glycine levels in the cell are high.

It can be assumed that this limiting condition is met by astrocytes, which accumulate glycine and whose endfeet surround capillary endothelial cells, near the largest and ultimate source of glycine. Therefore, sarcosine heteroexchange, which is fully functional at the negative resting potential of astrocytes, could scale up ambient glycine locally, potentially at higher levels than “true” GlyT1 inhibitors, which are dependent on the diffusion of glycine from an extracellular source or leakage by Asc-1, a sodium-independent amino acid transporter expressed in glia (Ehmsen et al., 2016).

The cytoplasmic accumulation of sarcosine by uptake and heteroexchange should lead to a decrease in glycine release over time. However, sarcosine is both a product and precursor of glycine, as both amino acids can be interconverted by one-carbon metabolism enzymes (glycine-N-methyltransferase for sarcosine synthesis and sarcosine dehydrogenase for its transformation in glycine (Ducker and Rabinowitz, 2017; Pérez-Sala and Pajares, 2023)). Therefore, a sustained and steady release of glycine by heteroexchange is possible provided that sarcosine is converted to glycine by sarcosine dehydrogenase, a mitochondrial enzyme expressed in astrocytes (Pérez-Sala and Pajares, 2023).

These glycine-releasing properties, combined with competitive inhibition of glycine uptake, may contribute to the clinical effect of sarcosine used as adjuvant therapy to improve negative and positive symptoms in schizophrenic patients (Tsai et al., 2004; Lane et al., 2005; Curtis, 2019). In addition, it should be noted that altered one-carbon metabolism has been associated with Schizophrenia (Smythies et al., 1997; Krebs et al., 2009).

## 5. ACKNOWLEDGEMENTS

I thank Annick Ayon for her help in molecular biology and Stéphane Dieudonné for generous discussions and support. I thank the National Institute of Health and Medical Research (INSERM) and the National Centre for Scientific Research (CNRS) for their support.

